# Minicells from Highly Genome Reduced *Escherichia coli*: Cytoplasmic and Surface Expression of Recombinant Proteins and Incorporation in the Minicells

**DOI:** 10.1101/2021.06.16.448719

**Authors:** Hanna Yu, Andrei V Khokhlatchev, Claude Chew, Anuradha Illendula, Mark Conaway, Kelly Dryden, Denicar Lina Nascimento Fabris Maeda, Vignesh Rajasekaran, Mark Kester, Steven L. Zeichner

## Abstract

Minicells, small cells lacking a chromosome, produced by bacteria with mutated *min* genes, which control cell division septum placement, have many potential uses. Minicells have contributed to basic bacterial physiology studies and can enable new biotechnological applications, including drug delivery and vaccines. Genome-reduced (GR) bacteria are another informative area of investigation. Investigators identified that with even almost 30% of the *E. coli* genome deleted, the bacteria still live. In biotechnology and synthetic biology, GR bacteria offer certain advantages. With GR bacteria, more recombinant genes can be placed into GR chromosomes and fewer cell resources are devoted to purposes apart from biotechnological goals. Here, we show that these two technologies can be combined: *min* mutants can be made in GR *E. coli*. The *minCminD* mutant GR *E. coli* produce minicells that concentrate engineered recombinant proteins within these spherical delivery systems. We expressed recombinant GFP protein in the cytoplasm of GR bacteria and showed that it is concentrated within the minicells. We also expressed proteins on the surfaces of minicells made from GR bacteria using a recombinant Gram-negative AIDA-I autotransporter expression cassette. As some autotransporters, like AIDA-I, are concentrated at the bacterial poles, where minicells bud, and because the surface-to-volume ratio of the small minicells is higher than bacteria, recombinant proteins expressed on surfaces of the GR bacteria are concentrated on the minicells. Minicells made from GR bacteria can enable useful biotechnological innovations, such as drug delivery vehicles and vaccine immunogens.

## Introduction

Investigators have studied the minimum complement of genes required for a living bacteria. Some investigators have taken an additive, synthetic approach ^1–3^. Other investigators have taken a subtractive approach, in which they make serial deletions in the bacterial chromosome. The Tokyo Metropolitan University Group, for example ^4, 5^, showed that they could delete up to 29.7% of the *E. coli* genome and still have viable bacteria. However, the doubling time is about twice as long for the highly deleted strains than for the wild type and the highly deleted strains exhibit altered morphology. Such studies offer insights into the minimal complement of genes needed for a viable bacterial cell. GR bacteria also present useful biotechnological applications. Eliminating genes not essential for growth can minimize diversion of energy and substrates for biotechnologically non-productive purposes or make additional chromosome capacity available for engineering goals. Eliminating genes involved in formation of bacterial structures may enable more effective studies of remaining structures. If animals are exposed to antigen over-expressing bacteria with many fewer functional genes, they have the potential to be less reactogenic.

Minicells (reviewed in ^6^) were described more than 50 years ago ^7^. In *E. coli*, proteins of the Min system, MinC, MinD, and MinE control the placement of the Z-ring in the middle of the bacterial cell by preventing assembly of the FtsZ complex at locations other than the middle of the cell. In wild type cells, these proteins promote cell division into approximately equal size daughter cells, and prevent formation of daughter cells lacking a bacterial chromosome. In *min* mutant cells, the Z-ring can form not only in the middle of the cell about to undergo division, but also toward one of the poles. When the Z-ring forms close to a pole, cell division yields a large cell with copies of the bacterial chromosome and a minicell lacking the chromosome. Mother cells containing bacterial chromosomes continue to divide, enabling ongoing minicell production. While daughter minicells are incapable of further reproduction, they remain metabolically and biosynthetically active. Minicells can be produced in quantity, using differential centrifugation and filtration approaches ^8, 9^.

Minicells have proved useful, both for the study of basic biological processes and for biotechnological uses. Minicells have been helpful in the study of bacterial processes and machinery, such as the flagellum and Type III secretion systems ^10, 11^ since studies employing cryo-electron tomography are easier. Minicells also played an important part in studies of bacteriophage physiology ^12–14^. Biotechnological applications include the use of minicells to encapsulate drugs ^9^, and have been proposed as potential vaccine antigen delivery vehicles ^15, 16^.

It may be useful to place recombinant proteins on the surfaces and make bacterial derivatives with enriched concentrations of recombinant surface proteins. Gram-negative autotransporters (autodisplay proteins, Type V secretion systems) (reviewed in ^17–19^) enable the placement of large numbers of recombinant proteins on the bacterial surface. Autotransporters have three domains: an N-terminal signal sequence that helps mediate transfer of the protein across the inner membrane via a Sec translocon mechanism, a central passenger protein domain that includes the effector portion of the protein, and a C-terminal β-barrel domain that intercalates into the outer membrane to form a pore-like structure, aided by the β-barrel assembly machinery (Bam) complex ^20^. The passenger domain transits out to the extracellular environment through the pore of the β-barrel. The β-barrel may exhibit chaperonin-like activity, aiding in the correct formation of passenger protein tertiary structure during transit to the extracellular environment. Subject to some limitations, for example an intolerance for disulfide bonds and certain size limitations, DNA sequences encoding heterologous proteins can replace native passenger protein coding sequence, so that autotransporters can be used to place heterologous recombinant proteins on the surface of the bacteria, anchored into the outer membrane by the β-barrel. Recombinant autotransporters have been used to place a variety of biotechnologically useful molecules on the surfaces of bacteria, including enzymes, biosensors, and vaccine antigens. However, no clinically useful vaccines have yet been produced using autotransporter expression systems, perhaps because the first vaccine applications were attempted using live bacteria as vaccine vectors with attendant safety concerns ^17, 18^, or the antigens expressed on the surfaces of the bacteria were insufficiently immunogenic.

At least some autotransporters preferentially accumulate at the bacterial poles ^21^, so it may be anticipated that recombinant proteins expressed on the surfaces of bacteria by means of the autotransporters will be enriched in minicells, since the minicells bud off from the poles. Combining substantial genome reduction with expression of recombinant proteins on the bacterial surface using autotransporters in combination with budding of the minicells from the bacterial poles would be expected to additionally enrich the relative amounts of the recombinant proteins on the surface of the minicells compared to expression of a protein in the cytoplasm or on the surface of non-genome reduced whole bacterial cells.

While both genome-reduced bacteria and minicells hold significant promise for a variety of biotechnological purposes, to our knowledge, these two technologies have not been previously combined. Here, we show that the *minC* and *minD* genes of highly GR *E. coli* can be deleted and that the resulting mutant bacteria are viable and biotechnologically functional.

## Results and Discussion

### *Production of the* minC/minD *mutations in genome reduced* E. coli *strains*

To explore whether it is possible to mutate the *minC* and *minD* genes of genome reduced bacteria and then make minicells from those mutant bacteria we obtained the comprehensive collection of genome deletions produced by the Tokyo Metropolitan University Group ^4, 5^ from a derivative of *E. coli* MG1655. We focused on the following strains: ME 5000 (parental, non-deleted strain), ME 5010 (2.4% deleted), ME 5119 (15.8% deleted) and ME 5125 (29.7% deleted, the most deleted strain in the collection). These strains, while they are alive, grow with a significantly reduced doubling time (~40 min for the 29.7% deleted strain), and altered morphology. We used lambda Red recombineering ^22^ to delete the *minC* and *minD* genes. We screened for the presence of the inserted marker, and confirmed the mutation by colony PCR and sequencing. We then cured the strain of kanamycin resistance by recombination with a derivative of pCP20 ^22, 23^ and verified the removal of the kanamycin resistance gene phenotypically, confirmed removal with sequencing, and reconfirmed the minicell production phenotype as described below. To examine expression in the minicells made from the genome reduced bacteria, we used a plasmid, pMP2463 that expresses GFP, and we commissioned the synthesis of a plasmid, pRAIDA2 (FIG 1), that includes an AIDA-I-based autotransporter expression cassette ^24, 25^. In its parental form, pRAIDA2 has an influenza HA immunotag in the autotransporter expression cassette.

**FIG 1.**
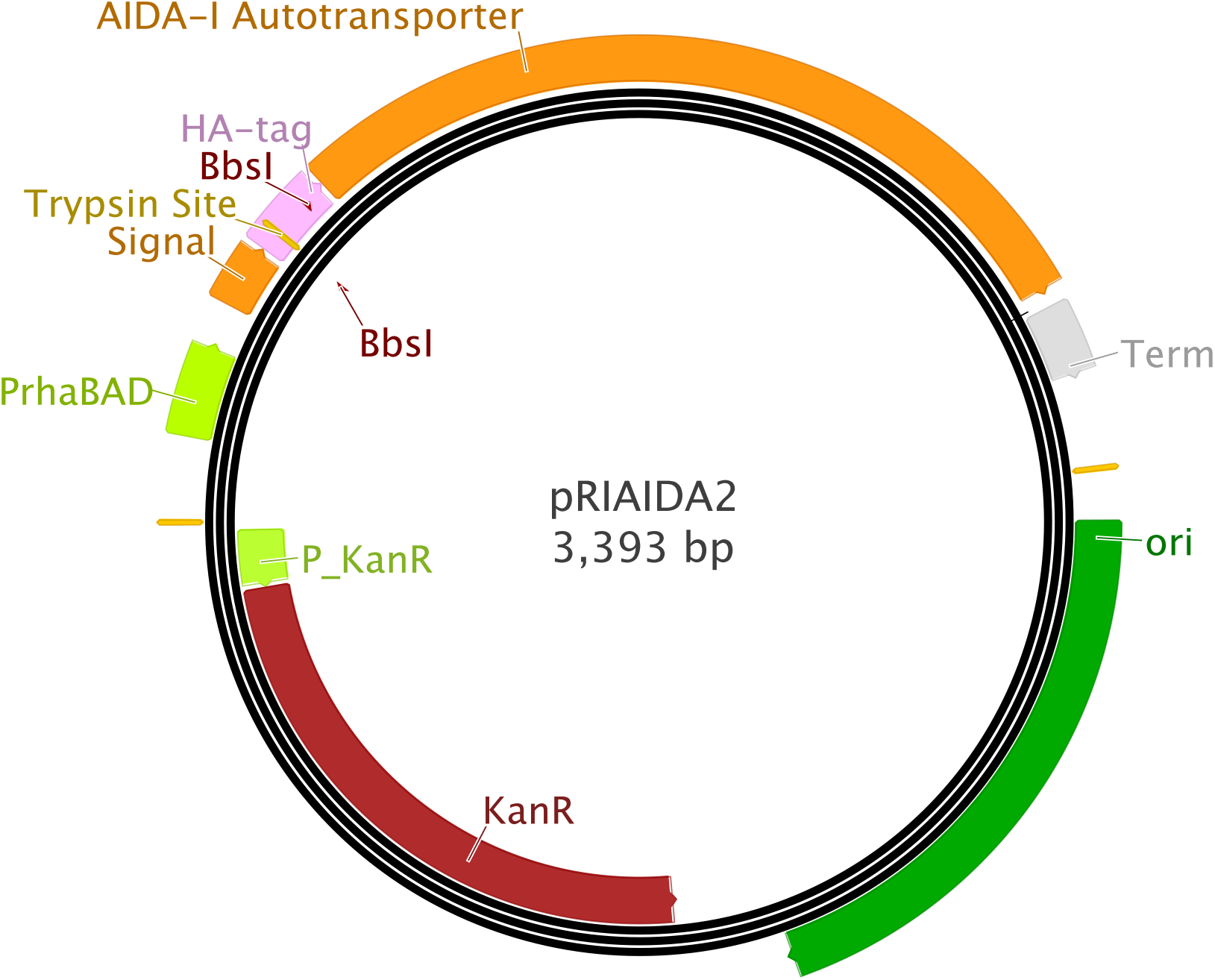
pRAIDA2 plasmid map. The plasmid has a high copy origin of replication, a kanamycin resistance gene, and an AIDA-I autotransporter under the control of a rhamnose inducible promoter. The parental version of pRAIDA2 enables the expression of an influenza HA immunotag on the surface of a bacterial cell

#### Characterization of the minicell production phenotype

We identified production of minicells by the wild type and genome reduced *E. coli* strains by cryo electron microscopy (cryoEM). FIG 2 shows the results of the cryoEM experiments. We found that we were able to produce minicells from the parental wild type strain (ME 5000), and from each of the highly deleted strains ME 5010 (2.4% deleted), ME 5119 (15.8% deleted) and ME 5125 (29.7% deleted), with the production of purified minicells after a differential centrifugation procedure. The cryoEM studies confirmed the presence of mostly minicells in the purified minicell preparation. In this report we will focus on a comparison of the non-deleted ME 5000 strain and the 29.7% deleted ME 5125 strain. Image cytometry also confirmed production of minicells from both the wild type and deleted strains (see Fig. 4 below).

**FIG 2.**
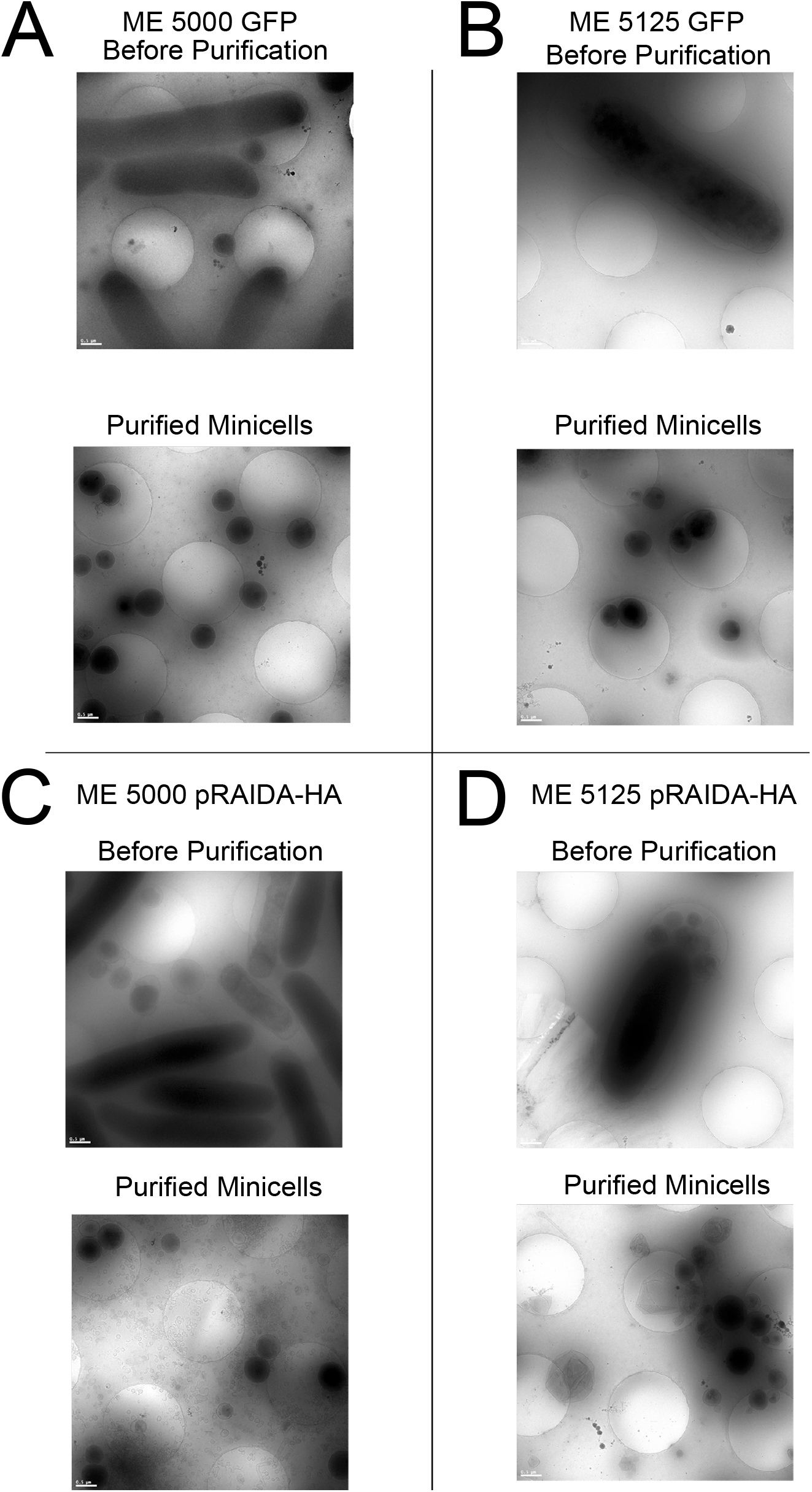
cryoEM images of cells and minicells from undeleted *E. coli* strain ME 5000 *minCminD* and highly deleted strain ME 5125 *minCminD*. The bacteria were transformed with either pMP2463, which expresses GFP, or pRAIDA2, which expressed the HA immunotag on the bacterial surface. A. *E. coli* strain ME 5000 *minCminD* transformed with the GFP-expressing plasmid pMP2463. B. Highly deleted *E. coli* strain ME 5125 *minCminD* transformed with the GFP-expressing plasmid pMP2463. C. *E. coli* strain ME 5000 transformed with the HA immunotag-expressing plasmid pRAIDA2. D. Highly deleted *E. coli* strain ME 5125 *minCminD* transformed with the HA immunotag-expressing plasmid pRAIDA2. The appearance of a large number of small, spherical minicells is evident in the cryoEM images for all strains and expression plasmid combinations tested.

#### Expressing recombinant proteins in the cytoplasm and on the surface of minicells made from genome reduced E. coli

Minicells made from genome reduced *E. coli* will likely have many uses, among the most important of these will be the production of minicells encapsulating biotechnologically useful proteins either in their cytoplasm or on their surfaces. To show that minicells made from genome reduced bacteria can contain recombinant proteins within their cytoplasm, we transformed the *minCminD* mutants of the wild type (ME5000) and the most highly genome reduced strain of *E. coli* in the TMUG collection (ME5125) ^4, 5^ with a GFP expressing plasmid, pMP2463 ^26^. We also transformed the undeleted ME 5000 and 29.7% deleted ME 5125 stains with *minCminD* mutations with pRAIDA2, the plasmid that expresses an HA immunotag via an AIDA-I autotransporter surface expression cassette under the control of a rhamnose-inducible promoter ^25^.

To determine relative amounts of recombinant protein in the minicells and parental cells, we conducted immunoblots on protein extracts made from the minicells and from the parental cells used to produce the minicells, interrogating the immunoblots with antibodies directed against either GFP expressed in the cytoplasm or the HA immunotag expressed on the surface of the cells, normalizing the signal attributable to the recombinant protein to a standard, the bacterial chaperone DNAK.

FIG 3 shows GFP and HA expression in the wild type and *minCminD* mutant ME5000 and genome reduced ME5125 *E. coli* strains as assessed by the immunoblots using antibodies against GFP and HA. We found that the GFP was present in both the parental cells and in minicells (FIG 3A), and that the normalized amount of GFP was enriched 8.7-fold in the minicells made from ME 5000, 2.3-fold in the minicells made from ME 5125 (FIG 3C). We also found that AIDA-I autotransporter-expressed HA immunotag was present in protein extracts made from the parental cells and the minicells (FIG 3B and FIG 3C). The HA immunotag placed into the minicells also appeared to be concentrated in the minicells. We found that HA was present in the minicells and was enriched 2.0 fold in the minicells made from ME 5000 and 1.9-fold in the ME5125 genome reduced bacteria. To analyze the data from the immunoblot experiment, we log-transformed the normalized densitometric data and compared the expression of GFP or HA in the minicells with the parental cells. The enrichments were statistically significant for the ME 5000 and ME 5125 cells expressing GFP or HA (p <0.01), by ANOVA analysis of the independently conducted immunoblot experiments.

**FIG 3.**
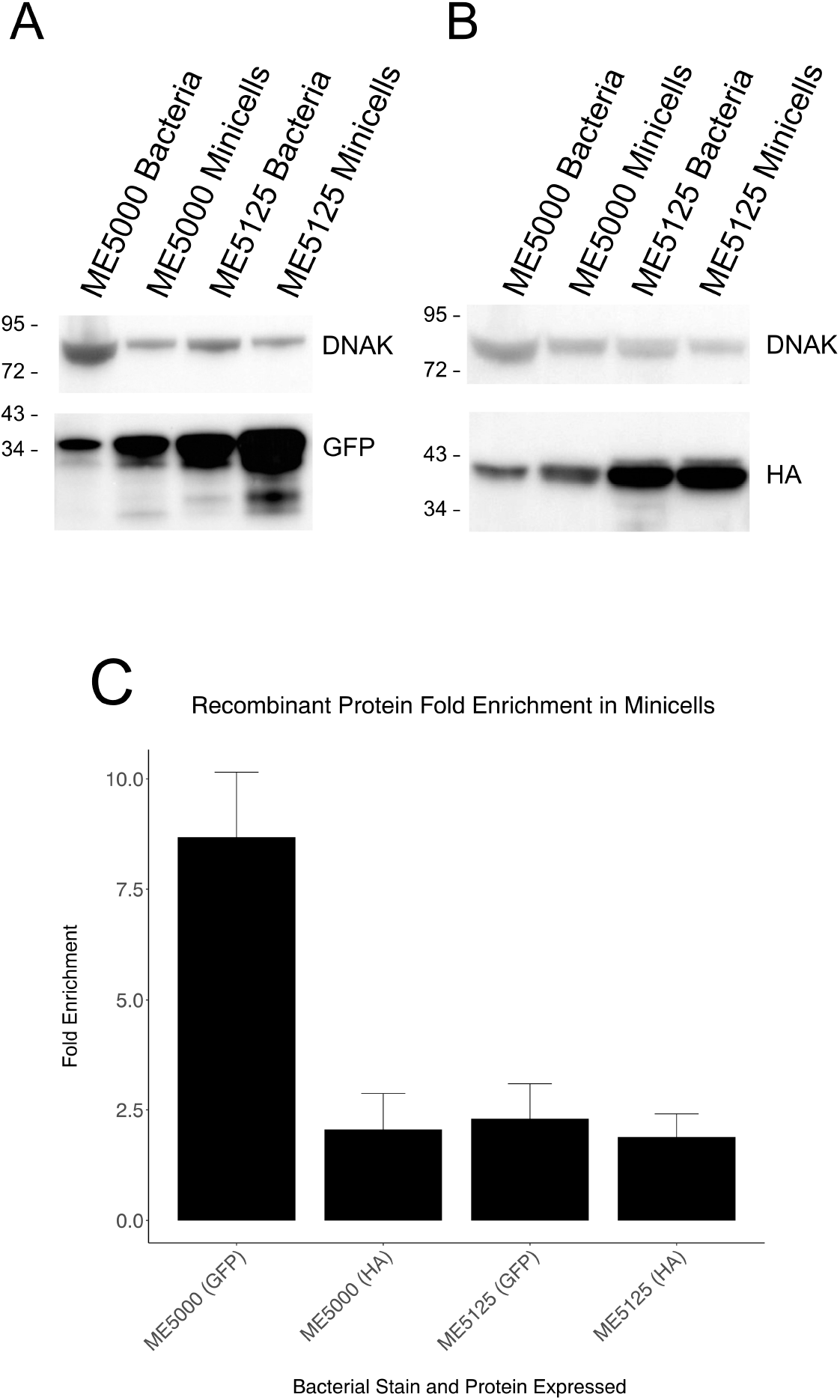
Immunoblots for recombinant proteins expressed in the cytoplasm (GFP) and on the surface (autotransporter-expressed HA immunotag) of non-highly genome reduced and highly genome-reduced bacteria, and minicells produced by the bacteria. A. Bacteria expressing GFP in the cytoplasm. Indicated lanes show the DNAK control normalization protein and the recombinant cytoplasmically expressed GFP in bacteria and minicells produced from the bacteria for the undeleted ME 5000 *minCminD* strain and the 29.7% deleted ME 5125 *minCminD* strain. B. Bacteria expressing HA via the AIDA-I autotransporter surface expression cassette. Indicated lanes show the DNAK control normalization protein and the HA immunotag in bacteria and minicells produced from the bacteria for the undeleted ME 5000 *minCminD* strain and the 29.7% deleted ME 5125 *minCminD* strain. C. Fold enrichment for the signal due to the indicated proteins, normalized to DNAK, in the minicell preparation compared to the bacteria.

#### Image cytometry analysis of non-genome reduced and genome reduce bacteria expressing GFP and HA

To further evaluate the expression of cytoplasmic- and surface-expressed recombinant proteins in non-genome reduced and genome reduced bacteria with *minCminD* mutations, we conducted a series of image cytometry studies. We imaged the bacteria, selecting images that the show bacteria with and without a budding phenotype (FIG 4). Here we show the images from the highly-deleted ME 5125 strain. The images show that GFP is expressed in the cytoplasm and that the GFP is also present in buds at the poles of the bacteria. The images further show that HA tends to be concentrated at the poles of the bacteria, as had been described previously for wild type *E. coli* ^21^. Some of the budding structures from the parent cell appear to show a strong concentration of the AIDA-I autotransporter surface-expressed recombinant HA in the buds.

To provide another quantitative evaluation of the distribution of the GFP and HA recombinant proteins in the minicells produced from the bacteria, we conducted an analysis of the minicells identified in the image cytometry studies. FIG 5 shows a selection of the image cytometry dot plots, with the gating strategies illustrated (FIG 5 A-F). We analyzed the signal intensity of the recombinant protein, either GFP or HA, per unit area of each identified particle as a metric of the amount of recombinant protein in the cells and their derived minicells. Elongated structures scatter more light than spherical structures, so particles gated as low scatter will preferentially represent minicells, while the particles gated as high scatter will preferentially represent the whole rod-shaped *E. coli*. The *minCminD* mutations block the proper placement of the cell division septum so, in addition to the minicells, cells mutant in *minCminD* produce elongated cells that retain the bacterial chromosomes, and this elongated phenotype can be appreciated in the images of FIG 4. More particles gated as high scatter can be appreciated in the *minCminD* mutants (panels D, E, F) in FIG 5. FIG 5 also shows that the GFP and HA recombinant proteins were concentrated in the minicells (compare for both the GFP-expressing cells, 5G, and HA-expressing cells, 5H, the signal intensity/area for ME 5125 Hi Scatter vs. ME 5125 Lo Scatter). The mean ratio of low scatter (minicells) to high scatter (parental cells) was 1.2 for the GFP-expressing ME 5000 cells, 1.3 for the GFP-expressing ME 5125 cells, 1.2 for the HA-expressing ME5000 cells, and 1.5 for the HA-expressing ME 5125 cells. The difference between the Signal Intensity/Area values between low and high scatter populations for both the GFP- and HA-expressing ME 5000 and ME 5125 cells was significant (p <0.001, twotailed t-test).

**FIG 4.**
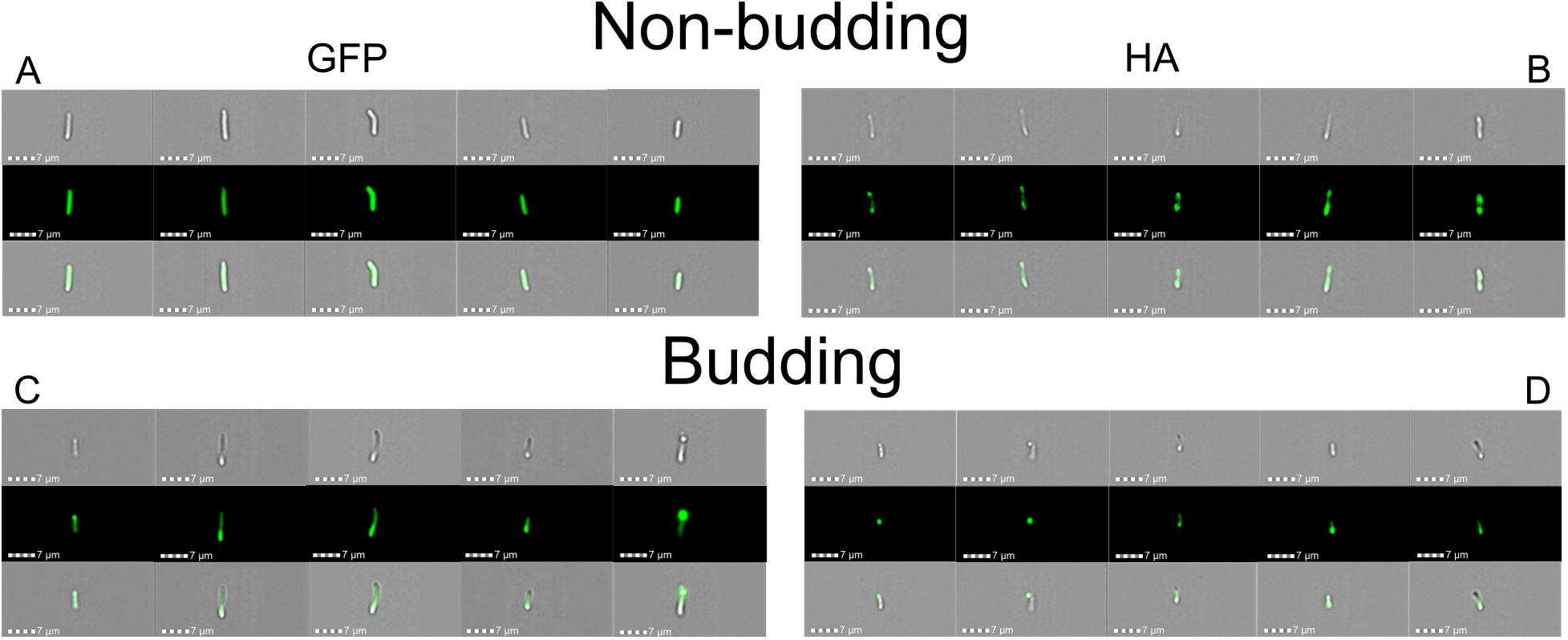
Examples of images and analysis of image cytometry data from ME 5125 *minCminD* expressing either the cytoplasmic protein GFP or HA expressed via the AIDA-I autotransporter, showing a non-budding or budding appearance. A. Non-budding cells expressing GFP. B. Non-budding cells expressing HA. C. Budding cells expressing GFP. D. Budding cells expressing HA via the AIDA-I autotransporter expression cassette. Of note, similarly to data previously described ^21^, for experiments involving the AIDA-I-expressed HA immunotag, the signal due to the autotransporter-expressed HA immunotag, concentrates at the poles of the bacteria (B), and then in these *minCminD* minicell producing mutant bacteria, appears to be further concentrated in the budding structures at the bacterial poles that are the precursors to minicells (D).

**FIG 5.**
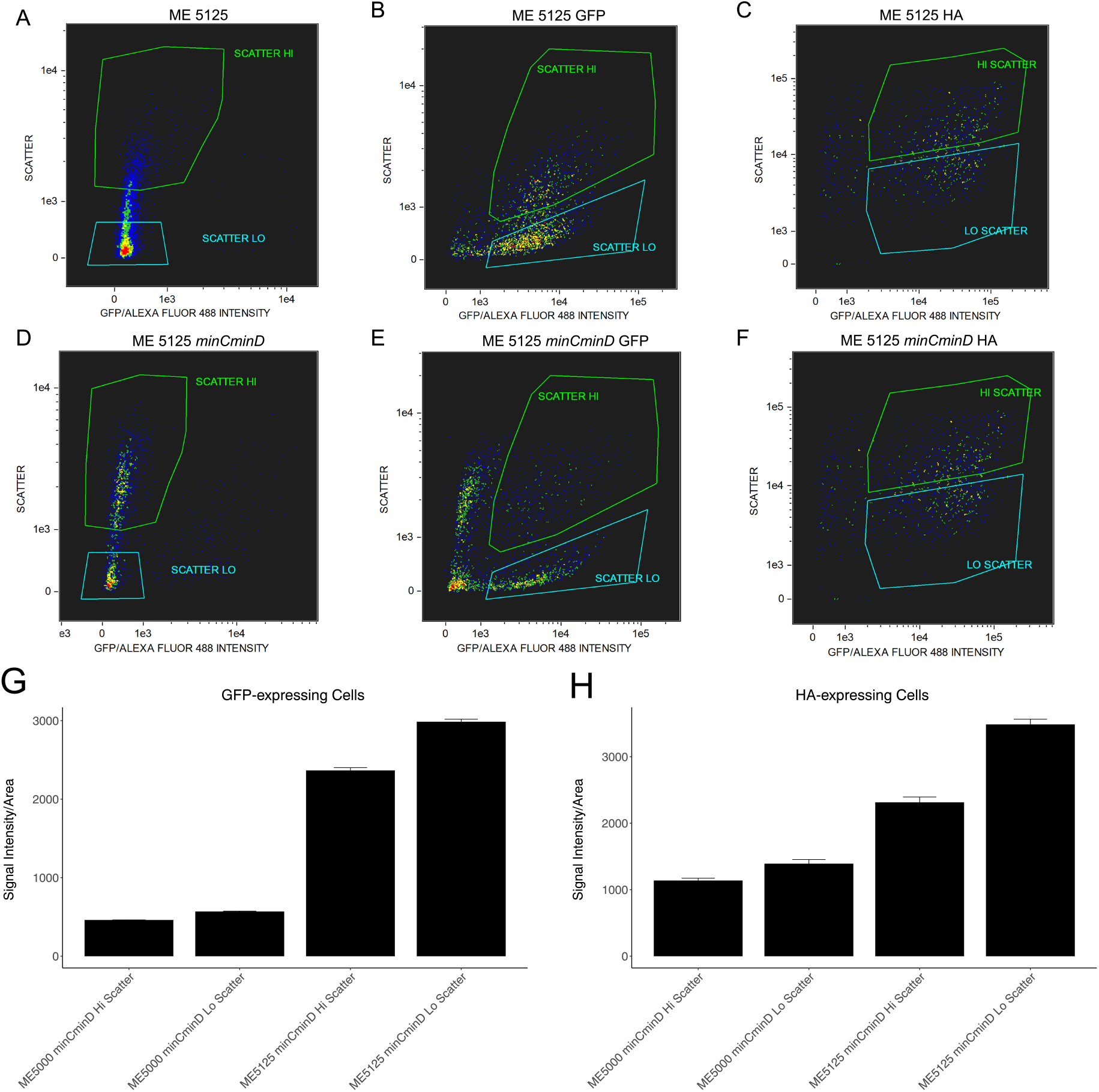
Image cytometry gating and analysis of differences in expression of the cytoplasmic protein, GFP, or the HA immunotag expressed via the AIDA-I autotransporter surface expression cassette. A. Image cytometry analysis of ME 5125 without an expressing plasmid. B. Image cytometry analysis of ME 5125 expressing GFP. C. Image cytometry analysis of ME 5125 expressing HA on the bacterial surface. D. Image cytometry analysis of ME 5125 *minCminD* without an expressing plasmid. E. Image flow analysis of ME 5125 *minCminD* expressing GFP. F. Image cytometry analysis of ME 5125 *minCminD* expressing HA on the bacterial surface. G. The ratio of the signal intensity from the fluorescence due to the recombinant protein GFP to the area of the gated particle (“Hi scatter” for the elongated bacteria, “Lo scatter” for the small, close-to-spherical mincells) in the preparation (arbitrary units), which contains both bacteria and derived minicells, for either the undeleted ME 5000 *minCminD* bacteria or the 29.7% deleted ME 5125 *minCminD* bacteria, as indicated. H. The ratio of the signal intensity from the fluorescence due to the recombinant HA protein to the area of the gated particle (“Hi scatter” for the elongated bacteria, “Lo scatter” for the small, close-to-spherical mincells) in the preparation (arbitrary units), which contains both bacteria and derived minicells, for either the undeleted ME 5000 *minCminD* bacteria or the 29.7% deleted ME 5125 *minCminD* bacteria, as indicated.

To confirm that the recombinant GFP protein and the HA immunotag expressed via the AIDA-I autotranporter are present in the minicells isolated from the *minCminD* mutants of the undeleted ME 5000 strain and the 29.7% genome-reduced ME 5125 strain, we conducted additional image cytometry experiments on minicells isolated from the bacteria using a differential centrifugation procedure. FIG 6 shows a selection of minicells isolated from the ME5000 and ME5125 strains expressing GFP or HA, demonstrating that the recombinant proteins are present in the minicells. In all cases, all minicells exhibited enhanced expression of GFP or HA.

**FIG 6.**
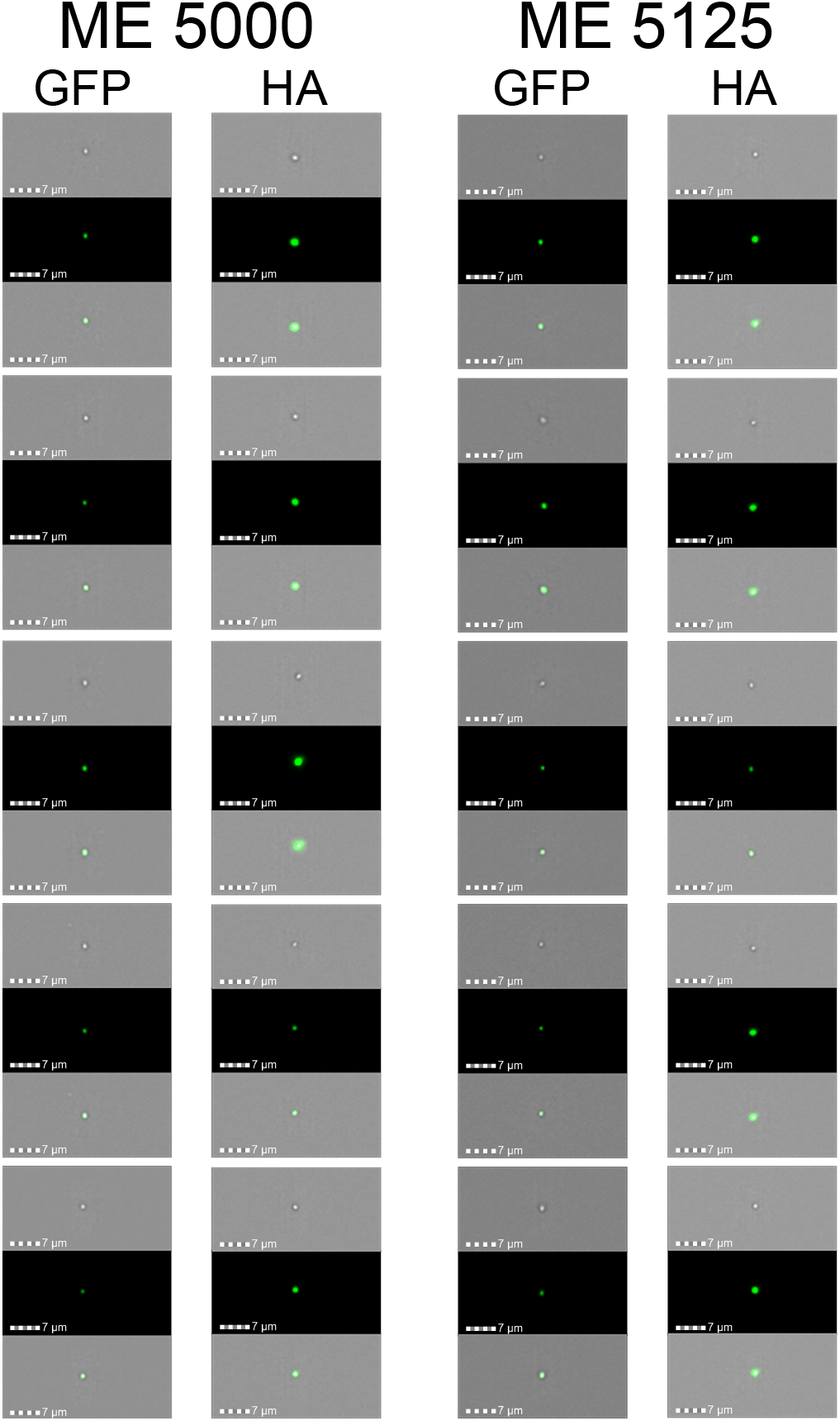
Examples image cytometry images of minicells isolated from undeleted ME 5000 *minCminD* or 29.7% deleted ME 5125 *minCminD*. The figure shows examples of five minicells analyzed by image cytometry from each bacterial strain, expressing either GFP or HA, as indicated. Each triple image shows a brightfield captured image (top), a fluorescent image of the minicell’s recombinant protein (middle), and a merged image (bottom).

Here we show that mutating *minC* and *minD* in GR *E. coli* strains produce minicells. Moreover, recombinant proteins can be successfully expressed in the genome-reduced cells, and concentrated in, and on, minicells derived from these genome-reduced *E. coli*. This demonstration can inform the use of minicells made from genome reduced *E. coli* in ways useful for basic investigations, bioindustry, and biomedicine. Additional advantages of a GR minicell may include improved cryo-EM structural studies due to fewer bacterial structures in the minicells as well as fewer genes that engage in metabolic activities not absolutely essential for metabolism, growth, and replication.

Bacteria are increasingly being metabolically engineered to more efficiently produce specific, industrially useful molecules (reviewed in ^27^). Unneeded bacterial metabolic pathways may divert substrates to non-useful pathways and/or produce unwanted side products. If minicells are used to package recombinant proteins or metabolic products made in engineered bacteria, using *minCminD* strains of genome reduced, engineered bacteria may be helpful.

Expressing recombinant proteins on the surfaces of bacteria using autotransporters can be a convenient way to express and purify proteins with limited solubility. Since translation and export to the bacterial surface is very tightly coupled, recombinant low solubility proteins are placed on the surface very soon after translation so they do not have an opportunity to form insoluble aggregates or inclusion bodies within the bacterial cell ^17–19^. Expressing recombinant proteins in genome reduced bacteria, followed by concentration in, or on, minicells may make this easier.

We demonstrated that both the cytoplasmically expressed protein GFP and AIDIA-I autotransporter-mediated HA immunotag were concentrated in the minicells. Concentration of the AIDIA-I autotransporter-mediated HA immunotag in the minicells would be expected because, for reasons yet to be established, AIDA-I concentrates at the poles of the bacilli, the sites from which the minicells bud. It is possible that the concentration factors that we observed are underestimates of the true value because, as evidenced by the DLS data of FIG 2 and the image cytometry data of FIG 4, the parental whole bacterial fractions contain both whole bacteria and minicells. At present, it is not possible, using differential centrifugation, to obtain a preparation of whole cells completely free of minicells; which could yield an underestimate of the amount of enrichment of the recombinant protein in the minicells compared to the parental cells. In addition, for the image cytometry studies, the gating strategies are to some degree arbitrary and cannot completely distinguish the minicells. It may be less obvious why the cytoplasmically-expressed GFP is concentrated in the minicells. However, in this case it may be helpful to consider that while the bacterial cytoplasm is sometimes thought of as being homogeneous, in fact it is not. The bacterial chromosome, and its associated proteins and structures, occupy a volume within the bacteria, so minicell cytoplasm is essentially derived from chromosome-free volume of the bacterial cytoplasm, where GFP may be present in higher abundance. Minicells may therefore be a means to enhance the relative amounts of bacterial non-chromosome-associated cytoplasmic proteins for bioindustrial processes.

One of the important uses proposed for minicells has been as antigen delivery vehicles for new vaccines ^15, 16^. No clinically approved vaccines have yet been produced using minicells, but it is plausible to hypothesize that a minicell with a surface-expressed vaccine antigen may be a more effective immunogen than a recombinant bacteria expressing the same antigen on its surface. An antigen expressed on the bacterial surface using a Gram-negative autotransporter would be more concentrated on the minicell. Minicells made from genome reduced bacteria would also include fewer host cell proteins. Minicells, because of their size and shape, may also transit across mucosal epithelial barriers more effectively and have more favorable interactions with immune system cells and so be useful for vaccine applications. We recently showed that killed whole cell *E. coli* vaccines that expressed antigens derived from the fusion peptides of either SARS-CoV-2 or porcine epidemic diarrhea virus (PEDV) can elicit good anamnestic responses and protect against disease in pigs in an experimental PEDV infection model. However, this approach did not elicit a strong neutralizing antibody response ^25^; an analogous minicell vaccine with an enhanced amount of antigen may yield improved immune responses. Approved killed whole cell vaccines have been produced at a cost of <1 US$ per dose ^28^, and we suggest that at-scale production of minicell vaccines should be not be substantially higher. If genome-reduced bacterial surface-expressed antigen minicell vaccines prove to have enhanced immunogenicity, such vaccines may prove useful globally.

In summary, we show here that minicells can be produced from highly genome-reduced *E. coli*, and that recombinant proteins are concentrated in the cytoplasm and on the surfaces of the minicells. Such minicells from GR bacteria may prove useful for a variety of biotechnological applications.

## Methods

### *Bacteria*, minC *and* minD *mutants*

The deleted bacterial strains with different percentages of the bacterial genome deleted were the kind gift of J. Kato ^4, 5^, Tokyo Metropolitan University, Tokyo, Japan. The National Bioresource Project, *E. coli* Strain Office, National Institute of Genetics, Japan, provided the *E. coli* strains used in this study, derivatives of MG1655, include ME 5000 (with 0% of the genome deleted – reference wild type), ME 5010 (2.4% deleted), ME 5119 (15.8% deleted) and ME 5125 (29.7% deleted). Strains were grown in Luria-Bertani (LB) media and on LB agar plates, with the appropriate antibiotics as needed.

### Electrocompetent cells and transformation with plasmid vectors

Bacteria were grown overnight at 37°C, shaking at 210 rpm in LB broth supplemented with selective antibiotic dependent strain specific resistance (streptomycin or ampicillin/chloramphenicol). New LB media cultures were inoculated from the overnight cultures and grown to log phase (OD_600_ ~0.4) with selective antibiotic. Due to their slower doubling times, the ME5125 strains required an additional day of growth in a new secondary inoculum between the primary inoculum and the third inoculum grown to log phase. The cells were centrifuged at 1,000 x g for 20 min at 4°C and washed with sterile ice-cold H2O + 10% glycerol. They were transformed by electroporation using the Gene Pulser Xcell electroporation system (Bio-Rad) with pRAIDA2, or pMP2463 ^26^. Electroporation was conducted in 0.1 cm electroporation cuvettes (Bio-Rad) with the following settings: 1800 V, 25 μF, 200 Ω ^29^. The electroporated cells were immediately transferred to 2 mL microfuge tubes with 1 mL of SOC media (Invitrogen), grown at 37°C, 80 rpm for 1 hour, and plated on LB agar plates containing 50 μg/mL kanamycin.

### Rhamnose induction

Bacteria transformed with pRAIDA2 were grown overnight in LB broth and 50 μg/mL kanamycin at 37°C, 210 rpm. The next day, new cultures were diluted to an OD_600_ of 0.1 in fresh LB broth and 50 μg/mL kanamycin, then incubated at 37°C, 210 rpm until the OD_600_ reached 0.4-0.7. GR ME5125 strains required an additional day of growth in a new secondary inoculum between the primary inoculum and the final inoculum grown to log phase (OD_600_ ~0.4). Bacterial cultures were induced with 5 mM of rhamnose for 2 hours. The OD_600_ of each culture was measured and aliquots of bacteria from each culture was saved for analysis by immunoblot. Afterwards, all cultures were centrifuged at 3,000 x g for 10 min at 4°C and pellets were stored in −20°C for ImageStream analysis.

We deleted the *minC* and *minD* genes using a modification of the Lambda Red recombination methodology ^22^. Briefly, the different *E. coli* strains were made electrocompetent by standard procedures ^30^. The cells were then transformed with pKD46, the plasmid having a thermosensitive origin of replication expressing the recombination machinery components (proteins Gam, Bet, and Exo) by electroporation using the Eppendorf Multiporator 4308, employing settings U = 2500 V, tau = 5 ms. Bacteria were then propagated at 30 C. We then produced a kanamycin resistance expression fragment for transformation and recombination with *minCminD* homology regions by PCR amplification using the pKD4 plasmid which includes the kanamycin resistance gene ^22^, as template. The PCR primers were: 5’-GTGACTTGCCTCAATATAATCCAGACTATAACATGCCTTATAGTCTTCGGAACATCATCGCGCGCTGGCGATGATTAATAGCTAATTGAGTAAGGCCAGGGTGTAGGCTGGAGCTGCTTC-3’ and 5’-CGCTGCGACGGCGTTCAGCAACAATAATCTGCAGCCGTTCTTTTGCAATGTTGGCTGTGTTTTTCTTCCGCGAGAGAAAGAAATCGAGTAATGCCATAACATGGGAATTAGCCATGGTCC-3’. The PCR reaction was performed using an Eppendorf Mastercycler Gradient 5331 (Eppendorf, Germany) with the following conditions: 95 C for 3 min; followed by 35 cycles of 95 C for 30 sec; 57.5 C for 30 sec; and 72 C for 2 min; followed by 72 C for 10 min; and then we held reactions at 4 C. PCR-amplified fragments were introduced into bacterial cells by electroporation using the Eppendorf Multiporator 4308, employing settings U = 2500 V, tau = 5 ms. After selection on kanamycin plates at 37 C, bacterial colonies were screened by PCR using primers 5’-GATTGAACAAGATGGATTGCACGC-3’ and 5’-CTCGTCAAGAAGGCGATAGAAGGC-3’, using the Eppendorf Mastercycler Gradient 5331 with the following conditions: 95 C for 2 min 30 sec; followed by 33 cycles of 95 C for 30 sec; 57 C for 30 sec; and 72 C for 1 min 40 sec; followed by 72 C for 10 min; and then samples were held at 4 C. PCR products were analyzed by agarose gel electrophoresis. The recombination event was verified by sequencing (Eurofins) to confirm replacement of *minC* and *minD* genes by the kanamycin resistance cassette. Bacteria were cured of the temperature sensitive origin of replication pKD46 plasmid by serial passage at 37C as described ^22^. Microscopic examination, dynamic light scattering, and image cytometry further confirmed the acquisition of the minicell production phenotype (see below). The *minCminD* mutant bacteria were then cured of kanamycin resistance by transformation with a FLP recombinase-expressing gentamycin-resistant modification of plasmid pCP20 ^23^ (kind gift of B. Doublet, INRA, Infectiologie Animale et Santé Publique, Nouzilly, France) ^22^ as modified ^31^. Individual colonies were picked and propagated overnight in the liquid culture at 43 C to induce recombination and loss of the pCP20-derivative plasmid. Restoration of a kanamycin-sensitive phenotype was verified by plating on kanamycin LB agar plates and sequenced to confirm removal of the kanamycin resistance gene. The minicell production phenotype was reconfirmed.

### Expression plasmids for evaluation of minicell production and expression of recombinant proteins

The green fluorescent protein (GFP)-expressing plasmid, pMP2463 ^26^, was obtained from addgene.com (https://www.addgene.org/107774/). We commissioned the synthesis of pRAIDA2 (FIG 1) (Genesys). pRAIDA2 includes an expression cassette based on the AIDA-I ^24^ autotransporter ^17, 32^ under the control of a rhamnose-inducible promoter, an origin of replication, and a kanamycin resistance gene ^25^. The parental version of pRAIDA2 expresses an influenza HA immunotag on a stuffer fragment within the expression cassette’s cloning site. The stuffer fragment is flanked by Bbs I, Type IIS restriction sites, to enable cloning of synthetic DNAs into the plasmid to enable surface expression of proteins of interest. The sequence of pRAIDA2 has been deposited into GenBank with accession number MW383928. Plasmids were prepared using Qiagen Plasmid Mini Prep kit, quantitated and assessed for quality spectrophotometrically.

### Expression plasmid transformation

To prepare competent cells, bacteria were grown overnight at 37 C, shaking, in LB broth. LB broth was inoculated from overnights and grown to log phase. Cells were made electrocompetent as described above and transformed via electroporation with pMP2463 or pRAIDA2. Electroporation was conducted in 0.1 cm electroporation cuvettes with the Gene Pulser Xcell electroporation system (Bio-Rad) and pulsed at the following settings: 1800 V, 25μF, and 200 Ω. Electroporated cells were immediately transferred to 10 ml tubes with 1mL of SOC media (Life Technologies), and grown at 37 C for 1 hour before plating on LB agar plates containing the appropriate antibiotic.

### Minicell purification and characterization

Minicells were produced as described ^33^ with modifications. A 3-5 ml miniprep culture of engineered bacteria was initiated from a glycerol stock or from a colony grown on fresh plate. The culture was grown at 37 C for 6-8 hours or overnight (in the case of ME5125 strain) and expanded to 50 ml culture growing overnight at 37 C. Next morning, this culture was used to start 1-2 liter culture which was grown 20-24 hours at 37 C with half of the usual antibiotic concentration. If required, the surface expression of the HA epitope was induced in the cells transformed with pRAIDA2 with rhamnose added to 0.5 mM added at the start of 1-liter minicells maxiprep.

Bacteria were separated from minicells by centrifugation 10 min at 4000 x g; supernatant was subjected to 10000 x g, 12 min centrifugation to sediment minicells. Minicell pellet was resuspended in 10-15 ml PBS and subjected to two subsequent 3000 x g, 10 min centrifugations to pellet residual bacteria. As the final step, to wash minicells from bacterial growth media, their volume was increased to 50 ml and minicells were spun down 10000 x g, 25 min at 15C. The final minicell pellet was resuspended in PBS and stored at −20C. The characteristics and quality of the minicell preparations were verified by dynamic light scattering and cryo-EM.

### Electron microscopy

The size and morphology of minicells were measured using a FEI Tecnai F20 (FEI, Hillsboro, OR) transmission electron microscope operating at 120 kV cryo-Electron microscopy (cryo-EM). The cryo-EM samples were prepared by a standard vitrification method. An aliquot of ~3 μl sample solution was applied onto a glow-discharged perforated carbon-coated grid, (2/1-3C C-Flat; Protochips, Raleigh, NC, USA) and the excess solution was blotted with filter paper. The samples were then quickly plunged into a reservoir of liquid ethane at −180 C. The vitrified samples were stored in liquid nitrogen and transferred to a Gatan 626 cryogenic sample holder (Gatan, Pleasantville, CA) and then maintained in the microscope at −180°C. All Images were recorded with a Gatan 4K x 4K pixel CCD camera under cryo-condition at a magnification of 9600X or 29,000X with a pixel size of 1.12nm or 0.37 nm, respectively, at the specimen level, and at a nominal defocus ranging from −1 to −3 μm. The unfiltered samples were recorded at 9600X, images were recorded with a 500nM magnification bar, filtered samples were recorded at 29,000X, and images were recorded with a t 100nM magnification bar.

### Immunoblots

Purified minicells and bacteria in the amount of 100-400 ug of protein were centrifuged for 15 minutes at 13-14000 x g at 10°C. Pellets were resuspended in NuPAGE 1x LDS electrophoretic sample buffer (Invitrogen) containing 3% 2-mercaptoethanol and heated at 100°C for 5 min. Samples in the amount 50-100 micrograms (GFP) or 200-400 ug (HA tag) were separated using NuPAGE 4 – 12% precast gels (Life Technologies) in SDS-containing buffer and transferred to Immobilon membrane (Bio-Rad). After 15 minute blocking in 1% casein blocker in TBS (ThermoFisher Scientific 37532), membranes were incubated with the primary antibody overnight at 4°C, washed and incubated with the horseradish peroxidase-conjugated secondary antibody. Primary and secondary antibodies were diluted in 0.05% TBS/Tween-20 containing 0.05% casein blocker. Protein bands were visualized using a chemiluminescence kit (ThermoScientific). Images were taken by ChemiXX6 G:box digital imaging system (Syngene) and quantified using GeneTools software (Syngene).

### Image Cytometry

Image cytometry assays were acquired on an ImagestreamX MKII (Luminex) using 488nm and 785nm Scatter lasers at 60X magnification to check (a) expression of green fluorescent protein in cytoplasm of bacteria transformed with pMP2463, (b) expression of HA-immunotag on outer membrane of bacteria transformed with pRAIDA2, and (c) cell morphology and size. Pellets stored in −20 C, after rhamnose induction, were thawed and formalin-fixed (1x HBSS + 0.2% formalin) and incubated at 37°C, 180 rpm for 1 hour. 5×107 formalin fixed and washed cells were blocked with 1x PBS + 10% FBS on ice for 20 min then washed and centrifuged at 800 x g with 1x PBS + 2% FBS for 5 min at 4°C. Cells were incubated with 1:200 dilution of HA tag monoclonal antibody (Invitrogen) for 30 minutes at 4°C. Cells were washed two times and centrifuged at 800 x g with 1x PBS + 2% FBS for 5 min at 4°C. Cells were incubated with 1:600 dilution of goat anti-mouse IgG Alexa488 (ThermoFisher) for 30 min at 4°C. After incubation, cells were washed two times and centrifuged at 800 x g with 1x PBS + 2% FBS for 5 min at 4°C then resuspended wash buffer and transferred into a 96 well flat bottom plate and stored at 4°C until analyzed by ImagestreamX MKII (Luminex).

The pellet of cells transformed with either pRAIDA2 (HA-expressing) or pMP2463 (GFP-expressing) were thawed post-rhamnose induction, formalin-fixed (1x HBSS + 0.2% formalin) and then incubated at 37°C, 180 rpm for 1 hour. Cells were washed with 1x PBS and centrifuged at 3,000 x g for 6 min at 4°C and then resuspended in 5 mL of 1x PBS before measuring the OD_600_nm. After calculations, 100 ul of 5×107 were aliquoted into the wells of a 96 well flat bottom plate and stored at 4°C until analyzed by ImagestreamX MKII (Luminex).

Eppendorf tubes containing minicells were spun down at 15,000 – 17,000 x g for 15 min at 4°C and supernatant was discarded. Pellets were resuspended in 1 mL of fixation solution (HBSS + 0.2% formalin) and incubated at 37°C, 180 rpm for 1 hour. 5×10^7^ formalin fixed minicells were washed and centrifuged at 10,000 x g for 12 min at 4°C before being blocked with 1x PBS + 10% FBS on ice for 20 min. The minicells were then washed and centrifuged at 800 x g with 1x PBS + 2% FBS for 4 min at 4°C. For minicells made from bacteria transformed with pRAIDA2, minicells were incubated with 1:200 dilution of anti-HA immunotag monoclonal antibody (Invitrogen) for 30 minutes at 4°C. Cells were washed two times and centrifuged at 800 x g with 1x PBS + 2% FBS for 5 min at 4°C. Cells were incubated with 1:600 dilution of goat anti-mouse IgG Alexa488 (ThermoFisher) for 30 min at 4°C. Post incubation, cells were washed two times and centrifuged at 800 x g with 1x PBS + 2% FBS for 5 min at 4°C then resuspended wash buffer and transferred into a 96 well flat bottom plate and stored at 4°C until analyzed by ImagestreamX MKII (Luminex).

Imagestream acquisition was set to collect 20,000 bacteria per sample. Objects were analyzed using IDEAS software 6.2.64.0. Focused bacteria were gated using the Gradient_RMS parameter for the brightfield channel. Single bacteria were then gated using the Aspect Ratio and Area parameters. Minicells were gated by Scatter and GFP/Alexa 488 signal intensity to distinguish whole rod, shaped bacteria from minicells.

### Statistical and data analysis

Data was analyzed with SAS 9.4 and R (v1.3.1093) with the Rstudio environment with included packages, and the tidyverse and stats packages, with visualization using ggplot2. The Western blot experiments were analyzed as randomized block experiments, using two-way ANOVA with no interaction. Data were transformed to the log scale to better meet the assumptions underlying ANOVA and to facilitate interpretations as fold change.

## Acknowledgements

Supported by the Wallace D. Coulter Foundation and by the Pendleton Pediatric Infectious Disease Laboratory, and institutional support for the University of Virginia Flow Cytometry Core Facility.

## Author contributions

H.Y., A.V.K., D.L.N.F.M, and V.R. designed and conducted the molecular biology, mutagenesis, bacterial transformation, recombinant protein expression, immunoblot, minicell production, and analysis experiments; C.C. conducted and analyzed image cytometry experiments; K.D. conducted and analyzed the cryo-EM experiments. M.C. provided statistical analyses; M.K. and S.L.Z. conceived, designed, and analyzed the experiments, drafted and revised the manuscript. All authors reviewed and approved the manuscript.

